# The impact of invader number on whole community invasions in biomethane-producing communities

**DOI:** 10.1101/2021.02.25.432953

**Authors:** Pawel Sierocinski, Jesica Soria Pascual, Daniel Padfield, Mike Salter, Angus Buckling

## Abstract

Microbes can invade as whole communities, but the ecology of whole community invasions are poorly understood. Here, we investigate how invader frequency affects the composition and function of invaded laboratory methanogenic communities. An invading community was equally successful at establishing itself in a resident community regardless of initial invader frequency, which varied between 0.01 and 10%. Invasion resulted in enhanced biogas production (to the level of the pure invading community), but only when invader frequency was 1% or greater. This inconsistency between invasion success and changes in function can be explained by a lower number of invading taxa (but not individuals) at lower initial invader frequencies, and an important functional role of the taxa that were absent. Our results highlight that whole community invasion ecology cannot simply be extrapolated from our understanding of single species invasions. Moreover, we show that methane production can be enhanced by invading poorly performing reactors with a better performing community at levels that may be practical in industrial settings.

## Introduction

Plant and animal invaders can play a major role in the structure and function of natural ecosystems [1–3]. Microbial populations and communities, like those of plants and animals, are also geographically structured [4–7], suggesting an important role of microbial invasions in their formation. We know relatively little about the cause and consequences of microbial invasions [8, 9], but there are clear parallels with the more extensive research on plant and animal invasion ecology [10–13]. In microbes for example, resident diversity typically reduces invasion success [14, 15], while invader population sizes [16–18] and disturbances [19, 20] may promote invasion. However, there are also likely to be major differences between microbial and macrobial invader dynamics. Notably, microbes often invade as whole communities, rather than as single species [21]. Leaves falling on the ground [22] or the release of sewage into the rivers [23] are examples of entire microbial assemblies arriving in a new ecosystem.

Recent theoretical and empirical work has started to identify conditions that affect the likelihood of whole community invasions [24, 25]. Resident diversity and productivity inhibit invasion [26, 27], while increasing invader diversity boosts invasion success (Rivett *et al*. 2018; Vila *et al*. 2019). Here, we focus on another theoretically important variable for both invading communities and populations: The number of invaders [17, 18, 28], or propagule size. A higher number of invaders can increase invasion success for a range of reasons. First, by reducing stochastic loss of invaders. Second, community size is likely to positively correlate with its species diversity [29, 30], meaning that a larger invading community is both more likely to contain a particularly invasive species, and show greater niche complementarity. The latter increases positive co-selection between community members [20, 26, 27]. Third, larger population sizes of invading species are also likely to result in invader populations adapting to new ecological conditions more rapidly [31].

While the number of invaders is expected to increase invasion success of single species and whole communities, the functional consequences of invasion may be less predictable for whole communities. Natural microbial communities are typically very diverse, meaning that stochastic loss of individual species within communities is increasingly likely with lower numbers of invaders. The rare taxa are more likely to be lost and there is growing evidence that rare taxa play a crucial role in community functions [20, 32, 33]. This means that invasions by lower numbers of invaders, even when successful, are less likely to be linked to key functional changes in the community than a similarly successful invasion initiated by a higher number of individuals. By contrast, a successful invasion by a single species might be expected to have similar consequences for community function regardless of initial number of invaders.

Here, we empirically investigate the relationships between invader numbers, invasion success, and functional consequences in methanogenic communities. Methane is an end product of community metabolism in these communities and therefore a useful proxy of community resource use efficiency (productivity). We use methanogenic communities because of their relevance to both biotechnology, where methane is playing an increasingly important role in the world’s renewable energy portfolio, and climate change in general, where melting permafrost may increase community mixing rates and impact methane release to the atmosphere [34]. Importantly, successful invasions are likely to result in increased methane production by the resultant community, as the higher methane producing communities have been shown to dominate when multiple communities are mixed together [27].

## Materials and Methods

### Experimental design

We used communities isolated from two anaerobic digester (AD) plants from the UK South West region. Before the experiment, we ran a pilot experiment for 4 weeks to measure methane production of each community. The stock communities were stored at 4°C for the duration of the experiment. We selected the community that produced the least methane of the two (Low Producer, LP) and cultivated 25 LP replicate fermenters using this community and 5 replicate fermenters of the community that produced more methane (High Producer; HP) in fermenters for two weeks to allow them to pre-adapt to experimental conditions. After two weeks, we invaded 20 of the LP replicates with either 10%, 1%, 0.1%, and 0.01% (by volume) of one of the HP replicates (5 replicates for each invasion percentage), while 5 LP replicates and 5 HP replicates were left undisturbed. The invaded LP communities and pure LP and HP communities were cultivated for an additional 6 weeks post-invasion, with gas production measured throughout and community composition determined at the beginning and end of the experiment.

### Cultivation and gas measurement

All replicates were grown in 500 mL bottles (Duran, 600mL total volume with headspace) using Automated Methane Potential Test System (AMPTS, Bioprocess Control Sweden AB)[27] to measure CO2-stripped biogas (Biogas) production. The pilot experiment and the pure LP and HP communities were started with 200 mL of the community. The 10% Invasion replicates were initiated with 180 mL of LP community and invaded with 20 mL of HP community after 2 weeks, while 1%, 0.1%, and 0.01% treatments were initiated with 198 mL of LP and invaded with 2 mL of either pure or HP, or HP diluted 10 or 100-fold with ddH2O, as appropriate. Each community was supplemented with 0.2 mL 1000x concentrated trace metal solution (1 g L^-1^ FeCl_2_. 4H_2_O, 0.5 g L^-1^ MnCl_2_. 4H_2_O, 0.3 g L^-1^ CoCl_2_. 4H_2_O, 0.2 g L^-1^ ZnCl_2_, 0.1 g L^-1^ NiSO_4_. 6H_2_O, 0.05 g L^-1^ Na_2_MoO_4_. 4H_2_O, 0.02 g L^-1^ H_3_BO_3_, 0.008 g L^-1^ Na_2_WO_4_. 2H_2_O, 0.006 g L^-1^ Na_2_SeO_3_. 5H_2_O, 0.002 g L^-1^ CuCl_2_. 2H_2_O) at the start of the experiment. In both the pilot and main experiments, all communities were fed weekly, starting on t_0_ with 2g of feed composed of 3.53% casein, 1.17% peptone, 1.17% albumen, 47.07% dextrin, and 47.07% sucrose (all compounds – Sigma), The feed was suspended in 18 mL sterile water.

### DNA extractions and sequencing

DNA was extracted and sequenced as described previously [33]. We extracted DNA using FastDNA™ SPIN Kit for Soil (MP). We confirmed the quality of extractions by electrophoresis on a 1% agarose gel, and we quantified it using dsDNA BR (Qubit). We constructed the 16S rRNA gene libraries using primers designed to amplify the V4 region [35] (Supplementary Table S1) and multiplexed. We generated amplicons using a high-fidelity polymerase (Kapa 2G Robust) and purified them using the Agencourt AMPure XP PCR purification system. They were quantified fluorometrically (Qubit, Life Technologies). We pooled the amplicons in equimolar concentrations based on Qubit results. We diluted the amplicon library pool to 2 nM in sodium hydroxide and transferred 5 μL into 995 μL HT1 (Illumina) to give a final concentration of 10 pM. We spiked 600 μL of the diluted library pool with 10% PhiX Control v3 on ice before loading into the Illumina MiSeq cartridge following the manufacturer’s instructions. The sequencing chemistry utilised was MiSeq Reagent Kit v2 (500 cycles) with run metrics of 250 cycles for each paired-end read using MiSeq Control Software 2.2.0 and RTA 1.17.28. One of the samples from HP treatment failed to sequence and is not included in the analysis.

### Data analysis

Sequencing data were analysed in R (v 3.3.2) using the packages *‘dada2’* [36], ‘*phyloseq’* [37], ‘*vegan*’ [38] and ‘*pairwiseAdonis*’ [39]. We used a full-stack workflow to estimate error rates, inferred and merged sequences, constructed a sequence table, removed chimeric sequences, and assigned taxonomy to each amplicon sequence variant (ASV) using the Greengenes database [40] following the published pipeline [36]. We estimated the phylogenetic tree using the R package ‘*phangorn*’, [41] where we constructed a neighbour-joining tree, and then fit a Generalized time-reversible with Gamma rate variation maximum likelihood tree using the neighbour-joining tree as a starting point. We used UniFrac distance (weighted and unweighted) as our measure of compositional dissimilarity between the communities (Lozupone & Knight 2005). In case of alpha diversity, we calculated species richness and evenness [43].

To establish if the invasions were successful, we needed to determine if organisms from the HP treatment managed to establish themselves in the invasion treatment. The majority of ASVs were shared between HP and LP samples (Fig. 1) making it difficult to discriminate relative community contributions using previous statistical methods [27]. Instead we subset the 16S sequencing data. For each invaded community, we determined the ASVs of attributable origin: those that were present in the endpoint of only one of the pure community treatments. ASVs found in LP community, but not in HP were attributed to LP and vice versa. ASVs present in both or neither pure community treatments were excluded. Invasion success was then estimated in two ways: as the proportion of ASV reads of HP origin in total reads (proxy for the fraction of successfully invaded individuals) or as a proportion of ASVs of HP origin in all ASVs (proxy for fraction of species that successfully invaded the community). Separate linear models were used to determine whether final invader number or invader diversity was altered by initial invader abundance, where invader abundance was treated as a categorical predictor. Model selection was done using likelihood ratio tests and post-hoc pairwise comparisons were between individual treatments.

**Figure 1.**
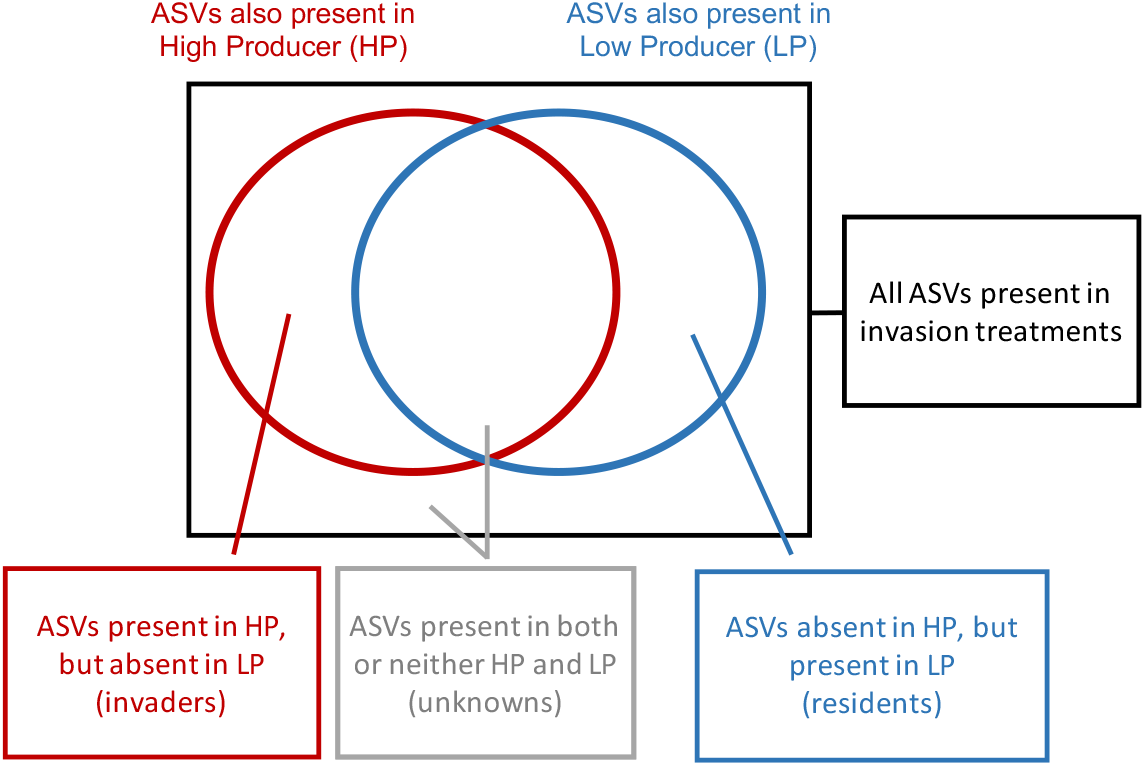
Schematic demonstrating how we attributed reads and ASVs to look at invader density and diversity. We checked if the ASVs found in the invasion treatments were also present in the LP and HP pure communities. ASVs found in one of the pure communities were attributed as originating from that pure community. ASVs found in both or neither pure communities were not attributed.

When looking at different community composition between treatments and pure LP and HP communities, we used all of the 16S sequencing data including all ASVs present in single treatments and pure LP and HP communities. We used a single permutational ANOVA to look at an overall difference in community composition between treatments, and used pairwise permutational ANOVA to see which treatments were driving any observed pattern. Differences in alpha diversity and evenness were tested using linear models, with number of ASVs or Pielou’s evenness as the response variable and treatment (invasion density or pure LP/HP community) as the predictor variable. Model selection and post-hoc comparisons were done as above. Similarly, gas production was compared using linear models, with gas production as a response variable and treatment (invasion density or pure LP/HP community) as the predictor variable. We used Dunnett’s Test to compare the invasion treatments to pure LP or pure HP gas production only, thus avoiding multiple comparisons between the invasion treatments. Model selection and post-hoc comparisons were done as above.

## Results

### Invasion Success

While only between 4 and 9% of all reads and between 40 and 47% of all ASVs in the invaded communities were attributable to only 1 community, the HP community appeared to successfully invade the LP community in all cases. The mean percentage of invading reads in total reads of attributable origin (Fig. 2A) varied between 8.5% and 60.3%. The mean percentage of invader ASVs in total ASVs of attributable origin (Fig. 2B) was between 16.3% and 46.6%. This means that even if all the non-attributed reads and ASVs originated from the LP community, as much as 0.7% and 7%, for total reads and numbers of ASVs, respectively, would still be attributable to the invading community (HP), much higher fraction than the lowest invader input of 0.01%. While the percentage of HP reads did not vary with the initial inoculum size of the invading community (Fig. 2B, F_1,18_ = 1.85, *P* = 0.19), the percentage of HP taxa was higher when initial inoculum size of the invading community was higher (F_1,18_ = 88.2, *P* < 0.001). There was an average of 90 invader ASVs in the 10% treatment, this number reduced to 53 ASVs in 1% treatment and 30 ASVs in 0.1% and 0.01% treatments.

**Figure 2.**
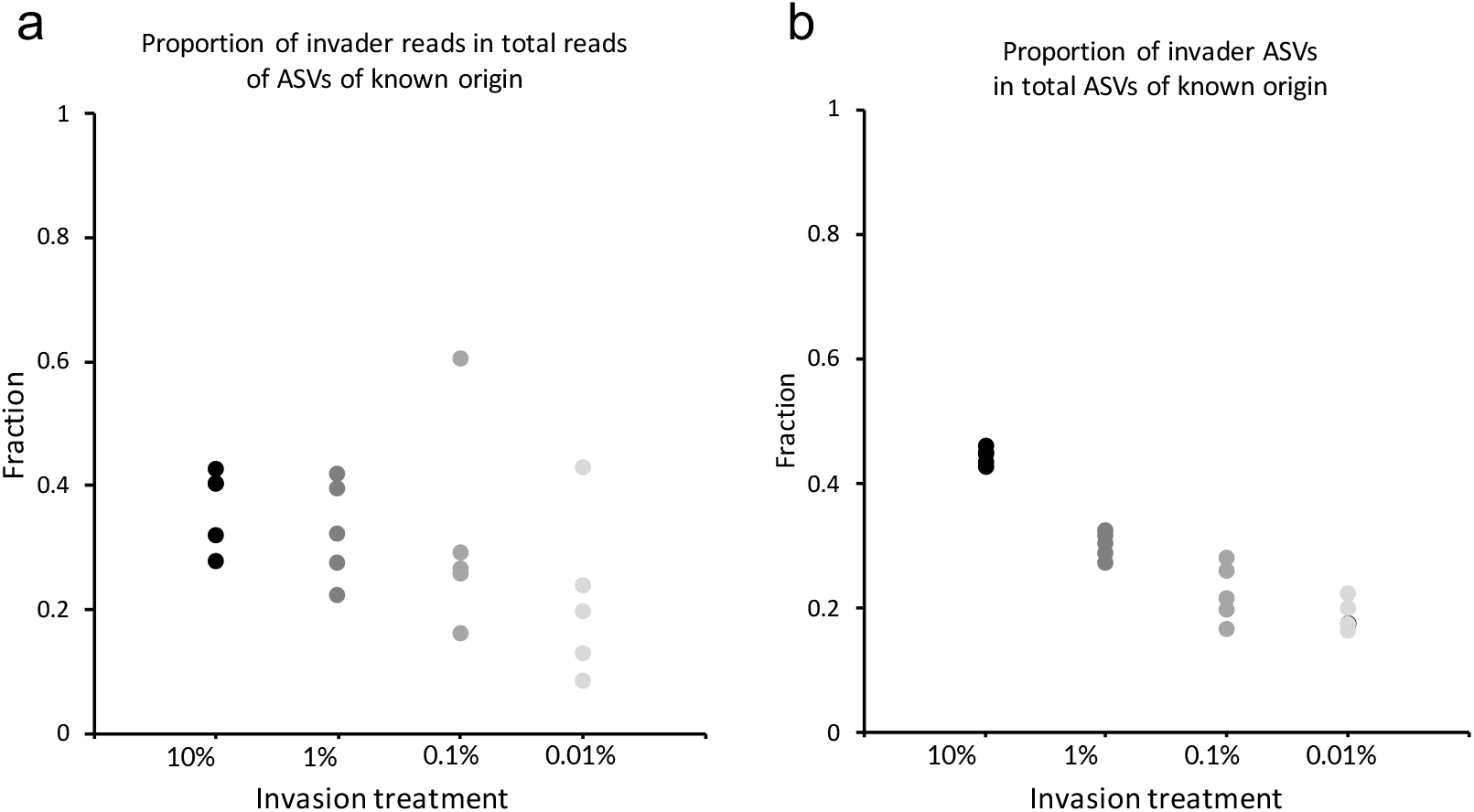
(A) Fraction of invader ASV reads in the total number of known-origin ASV reads (invader abundance); (B) Fraction invader ASVs in total known origin ASVs (invader diversity).

### Compositional changes following invasion

The composition of most of the invaded communities resembled the LP community when considering the abundances of all ASVs found in the samples (Weighted UniFrac): only the 10% (and HP) treatments differed in composition from the LP community (Fig. 3A, adonis2, F_5,23_ = 14.59 R^2^ = 0.76, *P =* 0.001, Pairwise adonis: *P* < 0.01). By contrast, all invaded communities were different from the LP community based on ASV presence and absence only (Unweighted UniFrac) (Fig. 3B, adonis2, F_5,23_ = 7.76, R^2^ = 0.62, *P* < 0.001. Pairwise adonis: *P* = 0.002). Note that communities changed considerably during propagation, with large differences in composition (using both metrics) largely because of ASV losses.

**Figure 3.**
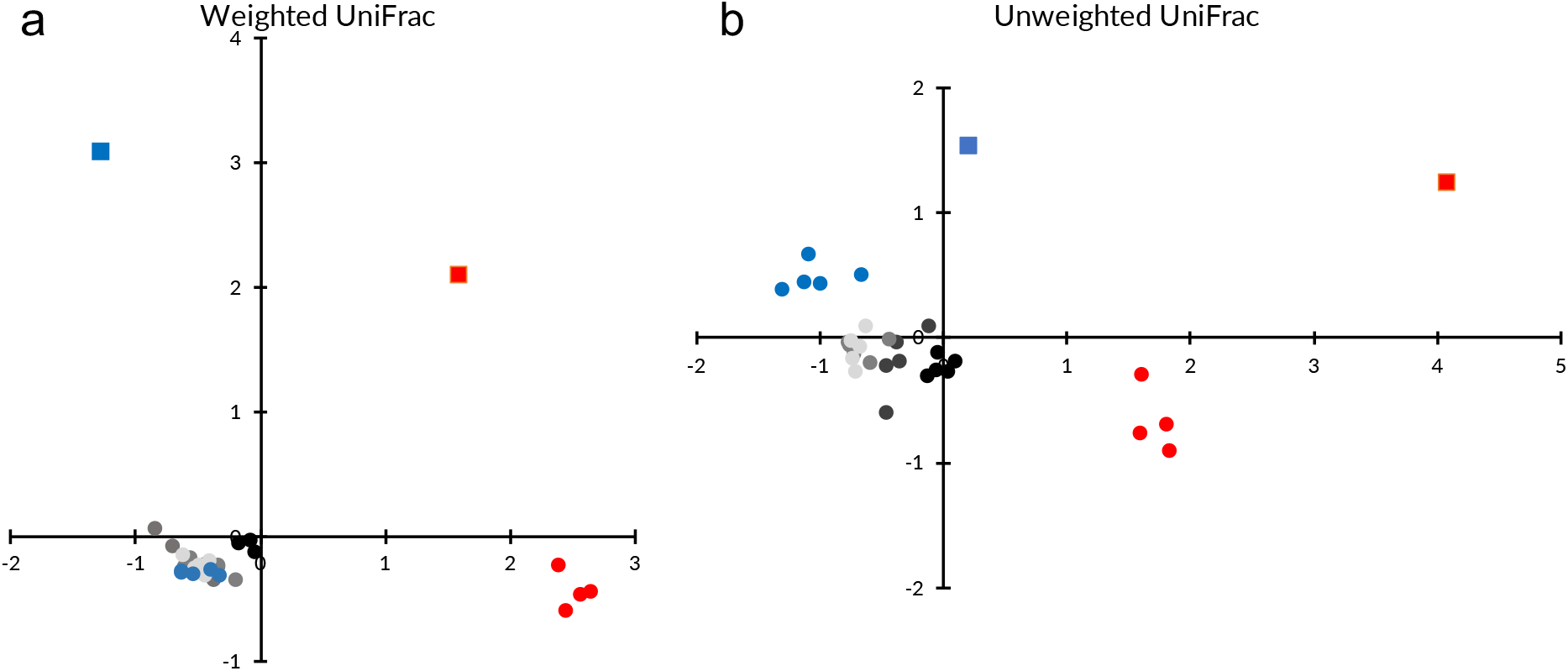
NMDS plot of: Weighted (A) and Unweighted (B) UniFrac of all the treatments. Red square: High producer ancestor; Blue square: Low producer ancestor; Red circles: High producer; Blue circles: Low producer, Black to light grey: Invasion treatments: in order from darkest to lightest: 10%, 1%, 0.1%, 0.01%

We also determined how invasion affected total within-community diversity. The HP community had significantly higher ASV richness than either the LP community or any of the invaded communities (Fig. 4A ANOVA, F_5,23_ = 24.5, *P* < 0.001, Tukey-Kramer HSD: *P* < 0.01), and there was a positive relationship between the richness of the invaded LP communities and the dose used in the invasion (F_1,27_ = 43.16, *P* < 0.001, R^2^ = 0.6). Species evenness (Pielou index) did not differ between the treatments (Fig. 4B, ANOVA, F_5,23_ = 0.3, *P* = 0.91).

**Figure 4.**
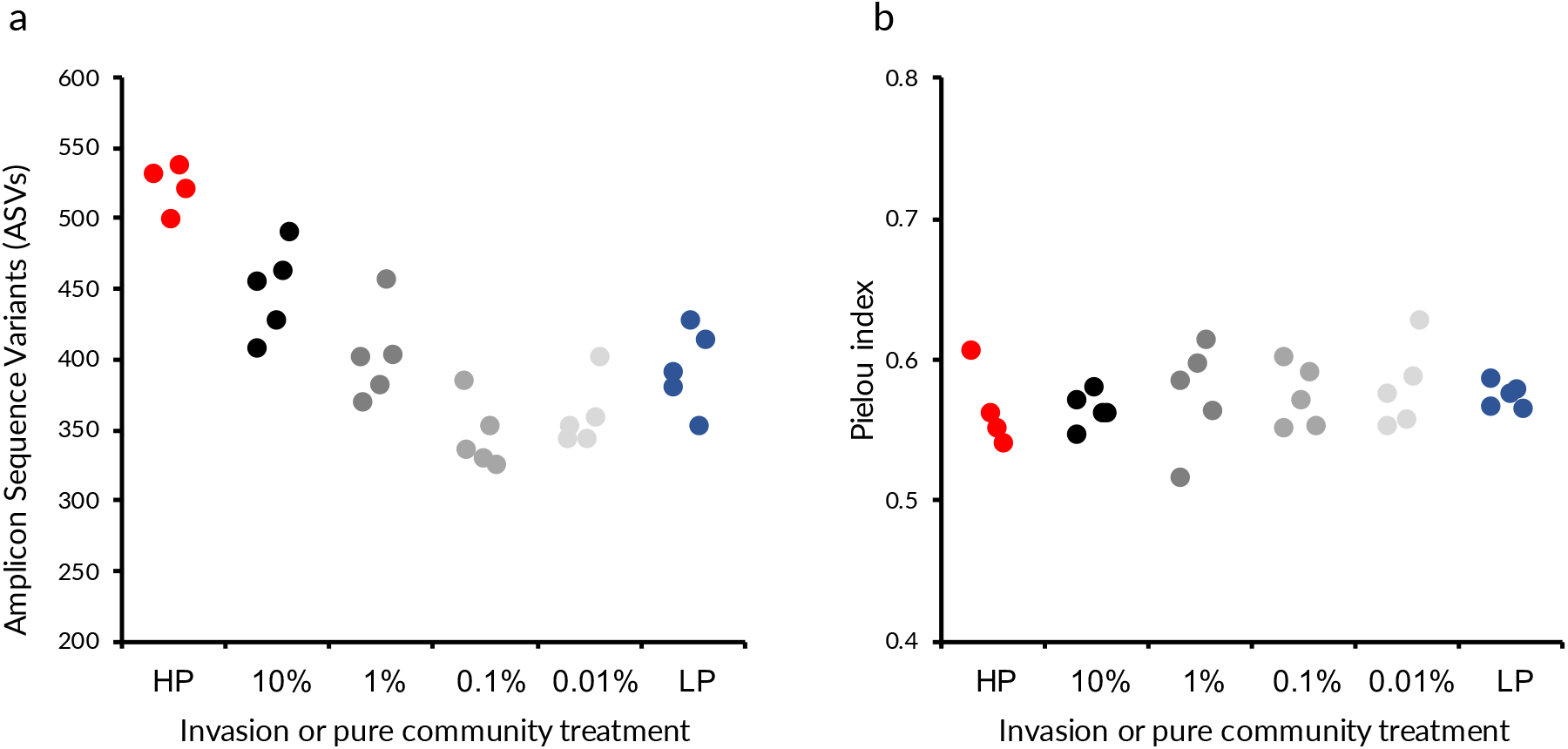
(A) Number of Amplicon Sequence Variants per sample (richness). (B) Pielou index (evenness).

### Gas production

Total gas production of LP communities increased with the log10 of invader inoculum size (Fig. 5B, F_1,23_ = 10.23, *P* = 0.003 R^2^ = 0.28). This enhancement of gas production resulted in the 10% and 1% invasion treatments showing gas production no different to the HP treatment, while the 0.1% and 0.01% invasion dose treatments produced significantly less biogas than HP (Fig. 5A, Dunnett Test with HP as control, *P* < 0.05 for significant differences) and conversely, 10% and 1% invasion treatments produce more gas than LP, while 0.1% and 0.01% treatments are no different from it (Fig. 5A, Dunnett Test with LP as control, *P* < 0.05 for significant differences)

**Figure 5.**
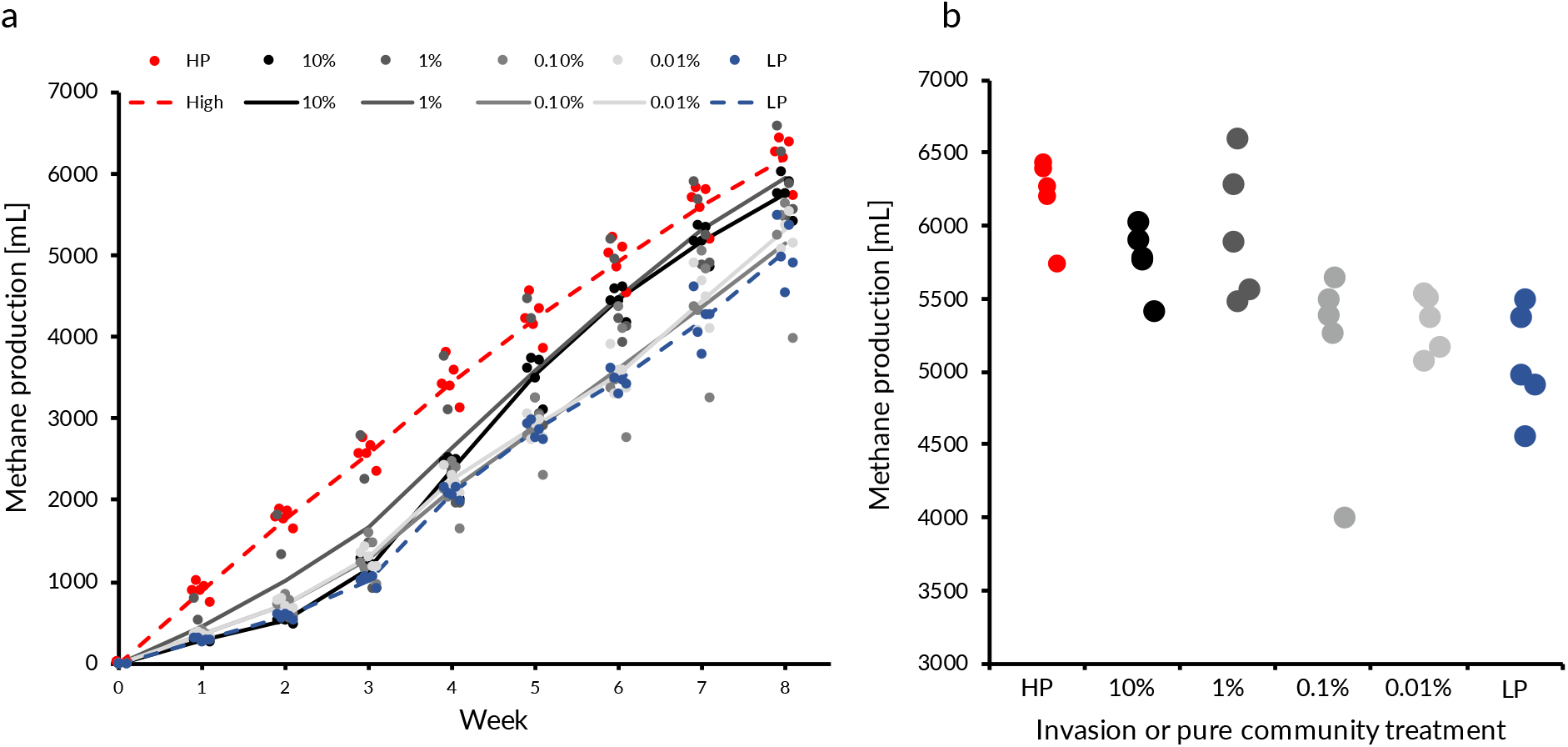
(A) Mean gas production over time. Points are individual replicates, while lines show treatment means. (B) Cumulative gas production at week 8.

### Species analysis

The communities were composed mainly of *Firmicutes* and *Bacteroidetes* phyla, with methane production driven by primarily acetoclastic methanogens in both communities. Given the apparent “tipping point” in gas production between 0.1 and 1% invasion, we determined if we could correlate this increase in function with particular taxa. We identified 44 ASVs (Supplemental material) that were present in the 10% and 1% but not the 0.1% or 0.01% treatment, and further refined these to 13 ASVs that were detectable in >50% of the 10% and 1% replicates. These 13 ASVs were mainly in the *Firmicutes* phylum, with species belonging to the *Ruminococcaceae* family being the most represented (4 ASVs). Aside from the Firmicutes, we found 2 abundant ASVs in the *Bacteroidetes* phylum from the *Porphyromonadaceae* family. One of those, a member of *Petrimonas* genus, was particularly numerous.

## Discussion

We investigated how the outcome of invasion was affected by the number of invading organisms in methanogenic communities. In contrast to our expectation, we found invasion success to be independent of the quantity of the invader used. Biogas production (a proxy for community productivity), was greater in the invaded communities (compared to the LP community), and increased with invader frequency. The relationship was stepped: gas production of the resident community was not altered by 0.01% and 0.1% invasion but resembled that of the invading community with 1 and 10% invader frequencies.

The disparity between the different measures of invasion success (number of individuals and number of taxa) is presumably because smaller number of invaders results in the loss of rare taxa. Either fewer taxa were initially inoculated into the resident community, or the reduced population size of some taxa resulted in stochastic loss during invasion. In contrast to recent work using synthetic communities [44], rare taxa did not appear to have contributed to the growth of the community as a consequence of co-selection between community members.

The loss of rare taxa with lower initial invader frequencies can also explain patterns of gas production. Increasing the volume of invading inoculum from 0.1 to 1%, led to approximately 40 more ASVs successfully invading and a marked increase in gas production. This is consistent with previous findings that the reduction in number of rare taxa through culture dilution reduced biogas production [33]. Note that the increase in gas production with invader frequency can’t be explained by key resources carried over from the media of the invading community, because there was a lag of approximately 4 weeks before gas production in these treatments started to increase.

While this work demonstrates that communities are capable of invading other communities from very rare, invading communities did not dominate the final community structure. Both theory and our previous experiments using methanogenic communities suggest that the most productive communities will dominate within a mixture of communities. However, in our previous study, unlike here, the communities were mixed at equal frequencies. The lack of invader dominating in low invasion treatments cannot be explained by invaders simply not having reached equilibrium conditions. The proportion of invader reads was the same regardless of invasion treatment, which means the communities stabilised before the endpoint of the experiment. This suggests there may have been a resident advantage through positive frequency or density dependence (a priority effect; e.g. see [45, 46]). This might arise because mutualistic interactions between residents are less disrupted when other community members are relatively rare.

Given the stepped increase in gas production with invader frequency, the taxa responsible for the increased gas production were presumably those present in the 10% and 1%, but not the 0.1% and 0.01%, invader treatments. Thirteen ASVs fitted that criterion and were present in more than half of the 10% and 1% invasion replicates. Only around seven of them could be identified to the family level. Four of the taxa present in majority of the treatments belonged to *Ruminococcaceae, two to Porphyromonadaceae* and one to *Syntrophomonadaceae* families, and are most likely involved in hydrolysis, protein metabolism and direct interspecies electron transfer [47–50]. While the first is not surprising, as the medium used in experiments is predominantly made from complex sugars, the other two point to interesting possibilities. Selection for organisms efficient at protein metabolism functions suggest a link between the increase of the community function and nutrient scavenging. The abundance of *Petrimonas* ASV from *Porphyromonadaceae* family could be linked to facilitating electron transfer between species and use of acetate in biogas production [50]. Given the dominance of acetoclastic methanogens in our bioreactors, this particular ASV is the most likely driver of increased biogas yield in the invaded communities

What is the relevance of our work to natural communities? Generalisations may be limited given that our laboratory conditions likely promoted invasion: Our communities were continually mixed [51] and experienced novel environments. Natural communities in stable environments are likely to be more locally adapted, promoting a resident advantage [46], and hence invasions will likely play a less important role in changes in community composition [52]. That said, our results are likely most relevant to communities experiencing environmental change, where an invading community is potentially better adapted to at least some local conditions than are the residents. An important implication is that while invasion may readily lead to restoration of a locally adapted community, this may not result in a concomitant restoration of function. methanogenic communities, which show much metabolic interdependence, are particularly prone to whole community invasions compared with other types of communities, such as soil [53], is unclear.

Our results suggest a potential novel approach to enhancing methane production in industrial settings, although further work is needed. We have previously established that seeding an Anaerobic Digester from a mixture of starting communities will, on average, result in improved gas production because the most productive communities tend to dominate. However, improving performance of existing reactors inevitably requires successful invasion by a smaller community than the resident. We show that invader frequencies as low as 0.01% can successfully invade. While the lower initial invader frequencies did not result in increased gas production, increasing absolute size of invader communities or (as would be the case when scaling up to industrial reactors) could overcome the problem associated with loss of rare taxa.

## Acknowldgements

This work was funded by BBSRC/Amur Energy IPA (BB/T002522/1).

## References

1. Montoya JM, Pimm SL, Solé R V. Ecological networks and their fragility. Nature. 2006. Nature Publishing Group., 442: 259–264

2. Wardle DA, Bardgett RD, Callaway RM, Van Der Putten WH. Terrestrial ecosystem responses to species gains and losses. Science (80-). 2011., 332: 1273–1277

3. O’Dowd DJ, Green PT, Lake PS. Invasional ‘meltdown’ on an oceanic island. Ecol Lett 2003; 6: 812–817.

4. Martiny JBH, Bohannan BJM, Brown JH, Colwell RK, Fuhrman JA, Green JL, et al. Microbial biogeography: Putting microorganisms on the map. Nat Rev Microbiol 2006; 4: 102–112.

5. DeLong EF, Preston CM, Mincer T, Rich V, Hallam SJ, Frigaard NU, et al. Community genomics among stratified microbial assemblages in the ocean’s interior. Science (80-) 2006; 311: 496–503.

6. Nemergut DDR, Schmidt SKS, Fukami T, O’Neill SP, Bilinski TM, Stanish LF, et al. Patterns and processes of microbial community assembly. Microbiol Mol Biol Rev 2013; 77: 342–356.

7. Whitaker RJ, Grogan DW, Taylor JW. Geographic barriers isolate endemic populations of hyperthermophilic archaea. Science (80-) 2003; 301: 976–978.

8. Mallon CA, Van Elsas JD, Salles JF. Microbial invasions: The process, patterns, and mechanisms. Trends Microbiol. 2015., 23: 719–729

9. Thakur MP, van der Putten WH, Cobben MMP, van Kleunen M, Geisen S. Microbial invasions in terrestrial ecosystems. Nat Rev Microbiol. 2019. Nature Publishing Group., 17: 621–631

10. Hall AR, Miller AD, Leggett HC, Roxburgh SH, Buckling A, Shea K. Diversity-disturbance relationships: Frequency and intensity interact. Biol Lett 2012; 8: 768–771.

11. Eisenhauer N, Schulz W, Scheu S, Jousset A. Niche dimensionality links biodiversity and invasibility of microbial communities. Funct Ecol 2013; 27: 282–288.

12. van Elsas JD, Chiurazzi M, Mallon CA, Elhottova D, Kristufek V, Salles JF, et al. Microbial diversity determines the invasion of soil by a bacterial pathogen. Proc Natl Acad Sci U S A 2012; 109: 1159–1164.

13. Lear L, Hesse E, Shea K, Buckling A. Disentangling the mechanisms underpinning disturbance-mediated invasion. Proc R Soc B Biol Sci 2020; 287: 20192415.

14. Levine JM, D’Antonio CM. Elton Revisited: A Review of Evidence Linking Diversity and Invasibility. Oikos 1999; 87: 15.

15. Hodgson DJD, Rainey PB, Buckling A. Mechanisms linking diversity, productivity and invasibility in experimental bacterial communities. 2002; 269: 2277–2283.

16. Lockwood JL, Cassey P, Blackburn T. The role of propagule pressure in explaining species invasions. Trends Ecol Evol 2005; 20: 223–228.

17. Ketola T, Saarinen K, Lindström L. Propagule pressure increase and phylogenetic diversity decrease community’s susceptibility to invasion. BMC Ecol 2017; 17: 15.

18. Acosta F, Zamor RM, Najar FZ, Roe BA, Hambright KD. Dynamics of an experimental microbial invasion. Proc Natl Acad Sci U S A 2015; 112: 11594–11599.

19. Lembrechts JJ, Pauchard A, Lenoir J, Nuñez MA, Geron C, Ven A, et al. Disturbance is the key to plant invasions in cold environments. Proc Natl Acad Sci U S A 2016; 113: 14061–14066.

20. Rivett DW, Jones ML, Ramoneda J, Mombrikotb SB, Ransome E, Bell T. Elevated success of multispecies bacterial invasions impacts community composition during ecological succession. Ecol Lett. 2018. John Wiley & Sons, Ltd., 21: 516–524

21. Rillig MC, Antonovics J, Caruso T, Lehmann A, Powell JR, Veresoglou SD, et al. Interchange of entire communities: Microbial community coalescence. Trends Ecol Evol. 2015., 30: 470–476

22. Koide K, Osono T, Takeda H. Colonization and lignin decomposition of Camellia japonica leaf litter by endophytic fungi. Mycoscience 2005; 46: 280–286.

23. Mansour I, Heppell CM, Ryo M, Rillig MC. Application of the microbial community coalescence concept to riverine networks. Biol Rev 2018; 93: 1832–1845.

24. Bellucci M, Bernet N, Harmand J, Godon J-J, Milferstedt K. Invasibility of resident biofilms by allochthonous communities in bioreactors. Water Res 2015; 81: 232–239.

25. Castledine M, Sierocinski P, Padfield D, Buckling A. Community coalescence: An eco-evolutionary perspective. Philos Trans R Soc B Biol Sci. 2020. The Royal Society., 375: 20190252

26. Tikhonov M. Community-level cohesion without cooperation. Elife 2016; 5: e15747.

27. Sierocinski P, Milferstedt K, Bayer F, Großkopf T, Alston M, Bastkowski S, et al. A Single Community Dominates Structure and Function of a Mixture of Multiple Methanogenic Communities. Curr Biol 2017; 27: 3390–3395.

28. Vila JCC, Jones ML, Patel M, Bell T, Rosindell J. Uncovering the rules of microbial community invasions. Nat Ecol Evol 2019; 3: 1162–1171.

29. Diamond JM, Mayr E. Species area relation for birds of the Solomon Archipelago. Proc Natl Acad Sci U S A 1976; 73: 262–266.

30. Bell T, Ager D, Song JI, Newman JA, Thompson IP, Lilley AK, et al. Larger islands house more bacterial taxa. Science (80-) 2005; 308: 1884.

31. Lachapelle J, Reid J, Colegrave N. Repeatability of adaptation in experimental populations of different sizes. Proc R Soc B Biol Sci 2015; 282: 20143033.

32. Jousset A, Bienhold C, Chatzinotas A, Gallien L, Gobet A, Kurm V, et al. Where less may be more: How the rare biosphere pulls ecosystems strings. ISME J. 2017. Nature Publishing Group., 11: 853–862

33. Sierocinski P, Bayer F, Yvon-Durocher G, Burdon M, Großkopf T, Alston M, et al. Biodiversity-function relationships in methanogenic communities. Mol Ecol 2018; 27.

34. Monteux S, Weedon JT, Blume-Werry G, Gavazov K, Jassey VEJ, Johansson M, et al. Long-term in situ permafrost thaw effects on bacterial communities and potential aerobic respiration. ISME J 2018; 12: 2129–2141.

35. Kozich JJ, Westcott SL, Baxter NT, Highlander SK, Schloss PD. Development of a dual-index sequencing strategy and curation pipeline for analyzing amplicon sequence data on the MiSeq Illumina sequencing platform. Appl Environ Microbiol 2013; 79: 5112–5120.

36. Callahan BJ, McMurdie PJ, Rosen MJ, Han AW, Johnson AJA, Holmes SP. DADA2: High-resolution sample inference from Illumina amplicon data. Nat Methods 2016; 13: 581–583.

37. McMurdie P, Holmes S. phyloseq: an R package for reproducible interactive analysis and graphics of microbiome census data. PLoS One 2013; 8(4): e61217.

38. Oksanen J, Kindt R, Legendre P. Vegan: Community ecology R package. R package version 2.2-1. 2007.

39. Martinez Arbizu P. pairwiseAdonis: Pairwise multilevel comparison using adonis. R package version 0.4. 2020.

40. DeSantis TZ, Hugenholtz P, Larsen N, Rojas M, Brodie EL, Keller K, et al. Greengenes, a chimera-checked 16S rRNA gene database and workbench compatible with ARB. Appl Environ Microbiol 2006; 72: 5069–5072.

41. Schliep KP. phangorn: Phylogenetic analysis in R. Bioinformatics 2011; 27: 592–593.

42. Lozupone C, Knight R. UniFrac: a new phylogenetic method for comparing microbial communities. Appl Environ Microbiol 2005; 71: 8228–8235.

43. Pielou EC. Species-diversity and pattern-diversity in the study of ecological succession. J Theor Biol 1966; 10: 370–383.

44. Goldford JE, Lu N, Bajić D, Estrela S, Tikhonov M, Sanchez-Gorostiaga A, et al. Emergent simplicity in microbial community assembly. Science (80-) 2018; 361: 469–474.

45. Connell JH, Slatyer RO. Mechanisms of Succession in Natural Communities and Their Role in Community Stability and Organization. Am Nat 1977; 111: 1119–1144.

46. Fukami T. Historical Contingency in Community Assembly: Integrating Niches, Species Pools, and Priority Effects. Annu Rev Ecol Evol Syst 2015; 46: 1–23.

47. Vartoukian SR, Palmer RM, Wade WG. The division ‘Synergistes’. Anaerobe. 2007. Academic Press., 13: 99–106

48. Grabowski A, Tindall BJ, Bardin V, Blanchet D, Jeanthon C. Petrimonas sulfuriphila gen. nov., sp. nov., a mesophilic fermentative bacterium isolated from a biodegraded oil reservoir. Int J Syst Evol Microbiol 2005; 55: 1113–1121.

49. Zhang J, Loh KC, Li W, Lim JW, Dai Y, Tong YW. Three-stage anaerobic digester for food waste. Appl Energy 2017; 194: 287–295.

50. Wang G, Li Q, Li Y, Xing Y, Yao G, Liu Y, et al. Redox-active biochar facilitates potential electron tranfer between syntrophic partners to enhance anaerobic digestion under high organic loading rate. Bioresour Technol 2020; 298: 122524.

51. Gravuer K, Scow KM. Invader-resident relatedness and soil management history shape patterns of invasion of compost microbial populations into agricultural soils. Appl Soil Ecol 2021; 158: 103795.

52. Ofiteru ID, Lunn M, Curtis TP, Wells GF, Criddle CS, Francis CA, et al. Combined niche and neutral effects in a microbial wastewater treatment community. Proc Natl Acad Sci U S A 2010; 107: 15345–15350.

53. Calderón K, Spor A, Breuil M-C, Bru D, Bizouard F, Violle C, et al. Effectiveness of ecological rescue for altered soil microbial communities and functions. ISME J 2017; 11: 272–283.

